# HAVEN: Hierarchical Attention for Viral protEin-based host iNference

**DOI:** 10.1101/2025.06.09.658367

**Authors:** Blessy Antony, Maryam Haghani, Adam Lauring, Anuj Karpatne, T. M. Murali

## Abstract

It is crucial to accurately predict hosts of viruses to understand and anticipate human infectious diseases that originate from animals. There is a lack of versatile models that handle out-of-distribution factors such as unseen hosts and viruses. We develop a machine learning model for predicting the host infected by a virus, given only the sequence of a protein encoded by the genome of that virus. Our approach, HAVEN, is the first to apply to multiple hosts and to generalize to unseen hosts and viruses. HAVEN is a transformer-based architecture coupled with hierarchical self-attention that can accept sequences of highly diverse lengths. We integrate HAVEN with a prototype-based few-shot learning classifier to predict rare classes. We demonstrate the accuracy, robustness, and generalizability of HAVEN through a comprehensive series of experiments. In particular, we show that HAVEN can achieve a median AUPRC of 0.67 while predicting common hosts. Moreover, HAVEN retains this AUPRC value even for rare hosts (median prevalence as low as 0.09%). Our model performs on par with state-of-the-art foundation models, which are 65 to 5, 000 times larger in size, and outperforms them in identifying hosts of SARS-CoV-2 variants of concern.

## 1 Introduction

Pathogens originating from animals have caused approximately 60% of infectious diseases in humans [1, 2]. There may be as many as 1.6M viruses that infect animals [3]. Thousands of these viruses could probably infect humans [4]. It is infeasible to experimentally study each virus to determine its host(s). Hence, we focus on the following classification problem: Given only the nucleotide or amino acid sequence of a viral genome or protein, predict which host the virus might infect. Accurate methods for this “ virus-host prediction” problem can help us better understand the origin of infectious diseases.

Virus-host prediction models must generalize in three ways: (i) Unseen virus: a model that generalizes to unseen viruses can help identify potential hosts for novel pathogens, aiding in early outbreak detection and response. (ii) Unseen hosts: the potential “ reservoir” host in nature of a newly-emerging virus may not be present in training datasets. (iii) Limited labeled data: there may be very few viral sequences available for hosts that are poorly studied due to geographical or logistical constraints.

Several studies have trained machine learning (ML) models to address this problem [5, 6, 7, 8, 9, 10, 11, 12, 13, 14, 15, 16, 17]. These methods have major limitations (Supplementary Note 1). Most models focus on only one viral species or family, e.g., influenza viruses, or a related group of proteins such as structural proteins of coronaviruses [5, 6]. Hence, they do not have the ability to generalize to unseen viruses or unseen hosts, e.g., given the sequence of an emerging virus, these methods are unlikely to predict its host in nature. Many existing approaches consider only binary classification [7, 8, 10, 12, 15]: does a given virus infect a specific host (usually, human) or not? Hence, these approaches will not be applicable to predicting the animal or reservoir host of a virus in nature. Some recent methods can predict multiple host labels across different taxonomic ranks. However, since these models were trained on organism-level links in virus-host networks, they cannot make accurate predictions given only the sequence of a viral protein [16, 17]. This limitation also applies to the network-based approaches that solved virus-host prediction as a link prediction problem in virus-host and virus-virus networks [9, 11, 13, 14].

To the best of our knowledge, ours is the first study to solve the virus-host prediction problem and address all the limitations. First, we did not restrict our focus to any protein family, which expands the scope and applicability of the model. Second, we considered multi-class prediction, i.e., we seek to predict the host species from a given viral sequence, encompassing both humans and a broad range of non-human hosts. Knowledge about the non-human host of a virus provides an understanding of its evolutionary trajectory and may inform risk mitigation strategies or countermeasures. Finally, we systematically demonstrated our method’s ability to meet all three generalizability challenges by integrating it into a few-shot learning (FSL) framework. Specifically, we accurately predict unseen and rare hosts. We also showed that we can predict seen and unseen hosts for unseen viruses. We believe that this is the first study to define and demonstrate all three desirable generalizability traits of a virus-host prediction model.

Our model, called ‘Hierarchical Attention for Viral protEin-based host iNference (HAVEN)’ used a pretrain-then-finetune paradigm based on the Bidirectional Encoder Representations from Transformers (BERT) [18] architecture. Rather than using one protein family, we pretrained HAVEN on the universe of all viral protein sequences, subject to restrictions on similarity that we describe later. To overcome the drawbacks of processing long protein sequences, we supplemented the BERT model with hierarchical self-attention [19]. We benchmarked the performance of HAVEN against standard machine learning classifiers, vision models adapted for sequence data, recurrent neural networks and their variants, and state-of-the-art protein language models (pLMs). When fine-tuned to predict host for a specific family of viruses, namely coronaviruses, HAVEN exhibited superior performance compared to state-of-the-art pLMs in accurately identifying hosts of SARS-CoV-2 variants.

## 2 Results

We defined the virus-host prediction problem as follows: given a protein sequence of any virus, predict the host infected by the corresponding virus. We developed ‘HAVEN’ to solve this classification problem. Starting from protein sequences from all viruses documented to infect vertebrates we curated multiple datasets (Section 3.1). We used different subsets of this universal dataset to evaluate HAVEN (Section 3.2) under various settings. We first assessed HAVEN’s ability to predict the most common hosts of non-Immunodeficiency viruses (non-IV dataset) (Section 3.3). Next, we fine-tuned HAVEN with FSL (Section 3.4) to predict unseen and rare hosts in the non-IV dataset. Finally, we gauged HAVEN’s performance in predicting hosts of unseen viruses using sequences of Immunodeficiency viruses (IV dataset).

We pre-trained HAVEN on 1.2 million viral protein sequences using masked language modeling (Section 3.3). Through this self-supervised learning process, our model learned to generate embeddings of protein sequences of any length. We fine-tuned this pre-trained model to predict virus-hosts in different scenarios.

### 2.1 Prediction of Common Classes in the non-IV Dataset

We first considered hosts with at least one percent prevalence in the non-IV dataset and fine-tuned HAVEN to predict five such “ common” hosts (Section 3.3). We compared the performance of HAVEN with the baseline models including, (i) standard ML classifiers: Logistic Regression (LR), Random Forest (RF), and Support Vector Machine (SVM), (ii) vision-based deep learning (DL) models: Convolution Neural Network (CNN), language-based DL models: Recurrent Neural Network (RNN), Long short-term memory (LSTM), and foundation pLMs: ProtT5, ProstT5, and ESM3.

We repeated all experiments five times with different splits of the dataset for training and testing. For each model, we measured the area under precision-recall curve (AUPRC) for each host to evaluate and compare the prediction performance (Section 3.3). We plotted the distribution of the macro-AUPRC (average of AUPRC for each host) scores over the five runs.

HAVEN with a median macro-AUPRC of 0.67 and inter-quartile range (IQR) of 0.02 performed significantly better than CNN (*p*-value= 3.97 × 10^−3^, Mann-Whitney U test), LSTM (3.97 × 10^−3^), ProstT5 (1.59 × 10^−2^). HAVEN was on par with ProtT5 (median macro-AUPRC=0.67; IQR=0.01) and ESM3 (0.65; 0.05) (Figure 2A). We then compared the models for each individual class (Figure 2B). All models have virtually similar performance for the dominant human class (prevalence=90.79%) with median AUPRC ≈ 0.99. HAVEN performed well in predicting the non-human classes despite their relative low prevalence in the labeled data: Pig (prevalence=4.04%; median AUPRC=0.60; IQR=0.09), Capybara (1.96%; 0.64; 0.16), Himalayan marmot (1.70%; 0.92; 0.06), and Red junglefowl (1.51%; 0.32; 0.04). ProtT5 and ESM3 also had comparable performance to HAVEN for some of the non-human classes.

**Figure 1:**
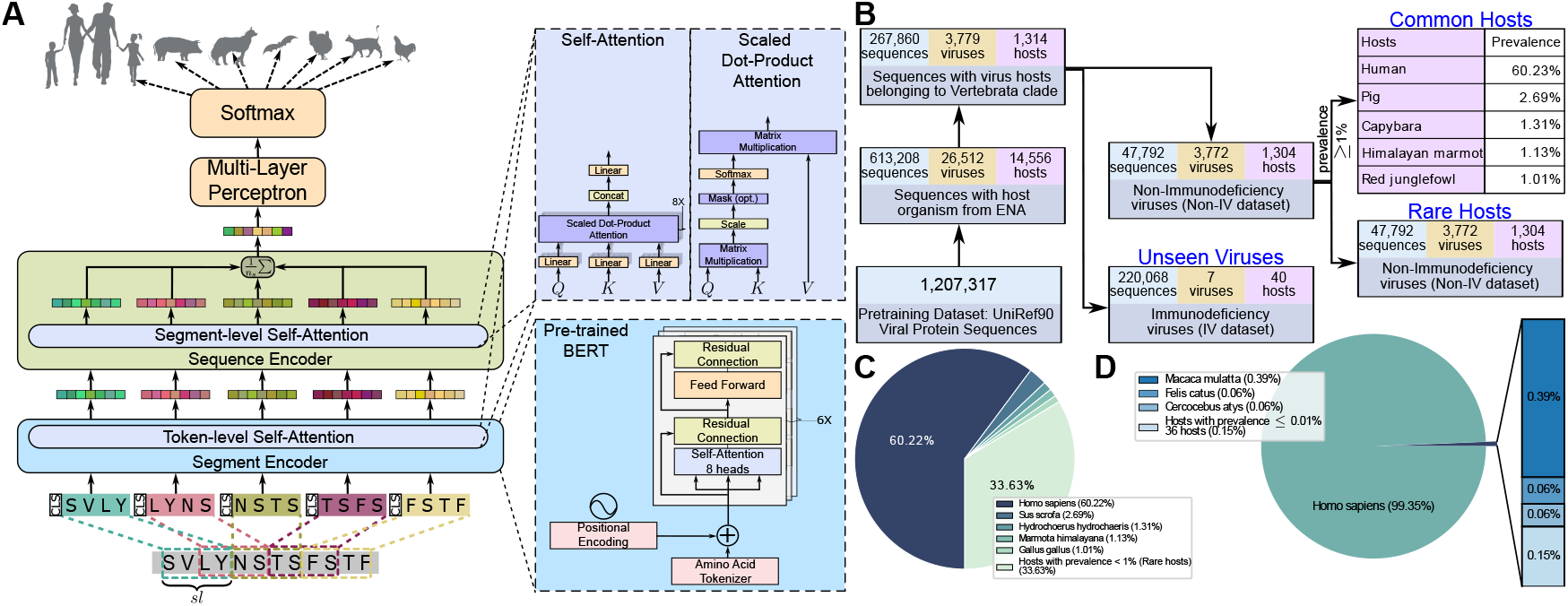
**(A)** HAVEN architecture. **(B)** Dataset Construction. Distribution of hosts in the **(C)** Non-IV dataset and **(D)** IV dataset. The Non-IV dataset consisted of five common hosts and 1, 299 rare hosts with prevalence less than 1%. The IV dataset comprised of 40 hosts with Homo sapiens (99.35%) dominating the dataset and 36 hosts with no more than 0.01% prevalence.

**Figure 2:**
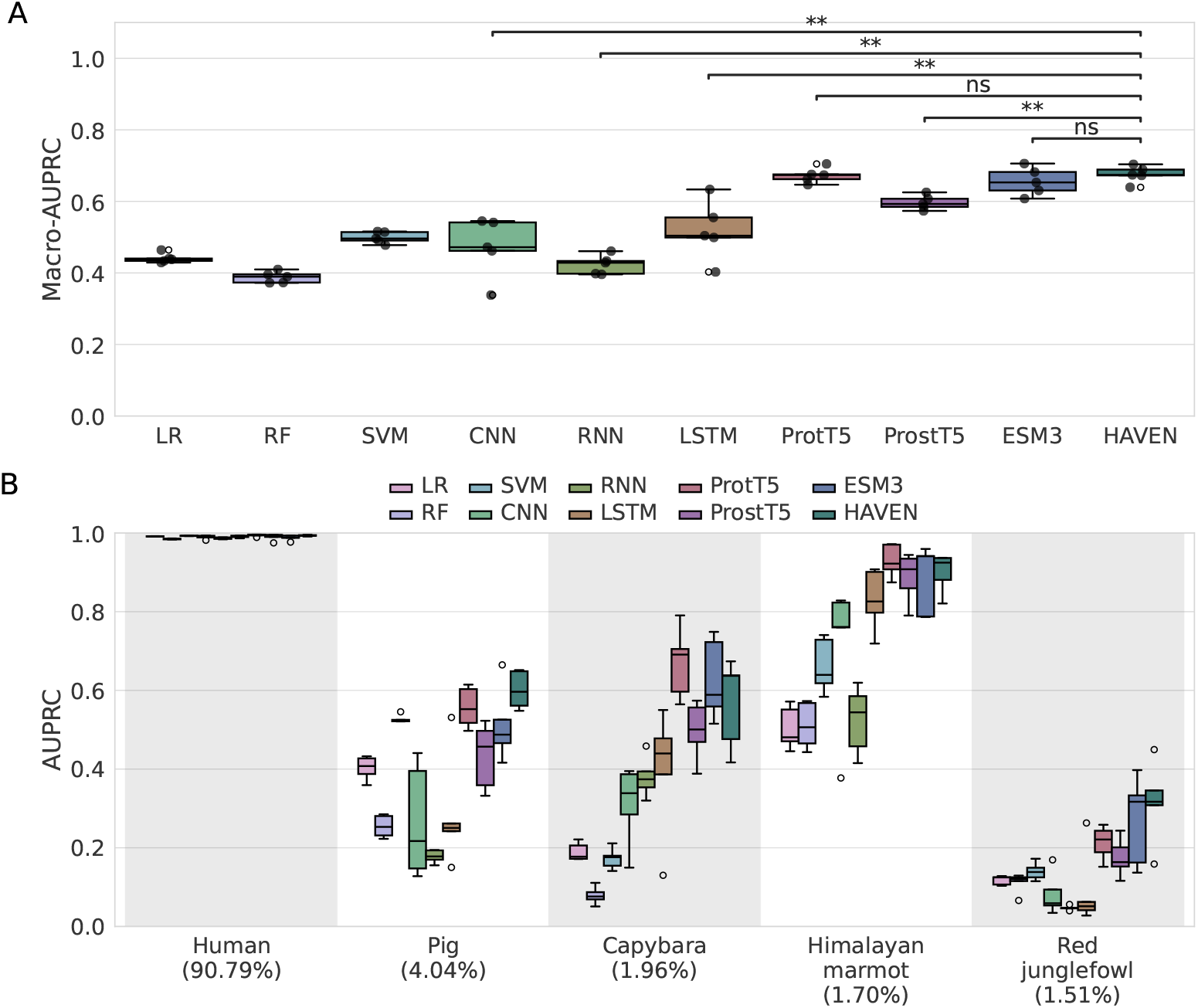
Distribution of the **(A)** macro-AUPRC and **(B)** classwise-AUPRC scores of all models in predicting the five common classes in the non-IV dataset. Abbreviations: LR, Logistic Regression; RF, Random Forest; SVM, Support Vector Machine; FNN, Fully-connected Neural Network; CNN, Convolution Neural Network; RNN, Recurrent Neural Network; LSTM, Long short-term memory.

### 2.2 Predicting Rare, Unseen Hosts

Encouraged by the performance of HAVEN for common classes, we measured its ability to predict rare classes using FSL in the non-IV dataset. Note that these hosts are unseen during the training done so far, by construction (Section 3.1). We trained and tested several prototypical network classifiers with varying *N*-way, *K*-shot configurations (Section 3.4, Supplementary Figures 5, 4, and 6). As expected, we observed a decrease in the class-wise AUPRC values as *N* increased from three to five in *N*-way, 5-shot classifiers and *K* decreased from five to one in the 3-way, *K*-shot classification. We selected the best performing 3-way, 5-shot classifier for subsequent analyses. Each data point in Figure 3 corresponds to a rare class from the test dataset selected in any one of the five iterations of FSL.

**Figure 3:**
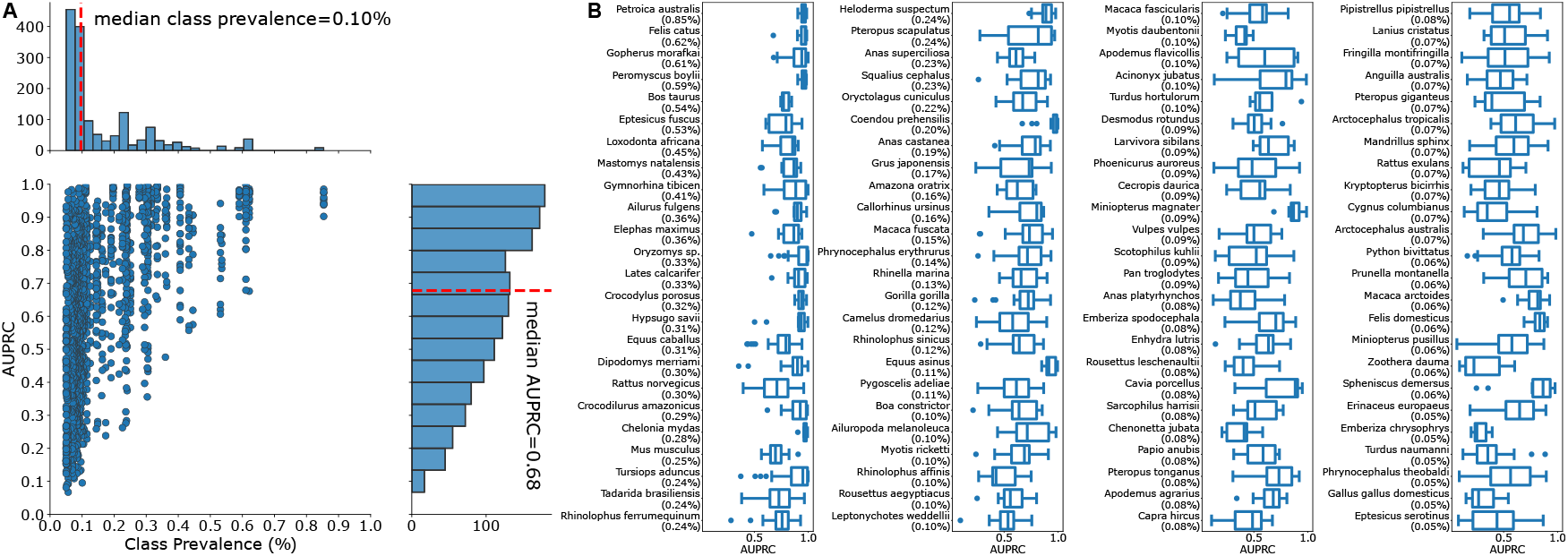
**(A)** Class-wise AUPRC of HAVEN fine-tuned using 3-way, 5-shot learning configuration to predict unseen rare host classes against the corresponding prevalence of the classes in the rare-host non-IV dataset. The X and Y axes marginals denote the distribution of the AUPRC scores and the class prevalences respectively. **(B)** Distribution of the class-wise AUPRC using HAVEN fine-tuned with 3-way, 5-shot FSL classifier to predict unseen rare host classes in the Non-IV dataset. The classes appear in the decreasing order of prevalence in the non-IV dataset.

Figure 3A shows the distribution of AUPRC scores of a 3-way, 5-shot FSL classifier for rare classes in the non-IV dataset plotted against class prevalence. The median AUPRC (across all iterations, epochs, and episodes) for each class ranged from 0.99 to 0.23 with a decreasing trend as the prevalence of rare classes changed from 1% to 0.05%. Notably, the median class prevalence was 0.10% (45 samples) but the median AUPRC was as high as 0.68, suggesting that HAVEN has the ability to make high-quality and reliable predictions of a virus’s host even when the number of sequences is very small.

Figure 3B shows the distribution of AUPRC values for each class across all episodes, epochs, and iterations. One of the four classes with the lowest prevalence of 0.05% (24 samples) was *Salmo salar* (Atlantic salmon). The AUPRC scores for this class ranged from 0.19 to 0.93 with median AUPRC of 0.51. This vast range in performance can be attributed to two factors namely, paucity of labeled data and the varying levels of complexity in distinguishing between the three randomly sampled rare classes in each episode of the 3-way, 5-shot FSL. Contrastingly, the AURPC scores of *Petroica australis* (South Island robin) class with the highest prevalence of 0.85% among the selected rare classes ranged from 0.90 to 0.98 with median value equal to 0.95. This higher performance for a comparatively more prevalent class reinforced the value of more labeled samples to learn about a host class.

### 2.3 Predicting Hosts of Unseen Viruses

We used the IV dataset (Section 3.1) to gauge the generalizability of the HAVEN model from the previous analysis in identifying hosts of unseen viruses. Our model did not encounter this dataset in any fine-tuning experiments done so far. We evaluated HAVEN under two different FSL prediction-only settings (Section 3.4). First, a 23-way, 5-shot classifier included the 23 unique hosts (four seen and 19 unseen) known to be infected by immunodeficiency viruses. For every sample in the IV dataset, the classifier computed a probability for each of the 23 classes. We also placed each sequence in a category based on the rank (out of 23) of the probability predicted for the true class (correct prediction); the categories were ‘Top rank’ (rank=one), ‘Top 3 ranks’ (3 ≤ rank < 1), ‘Top 5 ranks’ (5 ≤ rank < 3), ‘Top 10 ranks’ (10 ≤ rank < 5), or ‘Rank > 10’ (Figure 4B). The proportion of correct predictions for sequences from the dominant *Homo sapiens* (Human) class (218, 631; prevalence=99.347%) was 45.8%. We recognize that these accuracies can be significantly improved. It is likely that using only five sequences to compute the prototype for the human class does not capture the diversity of HIV sequences.

**Figure 4:**
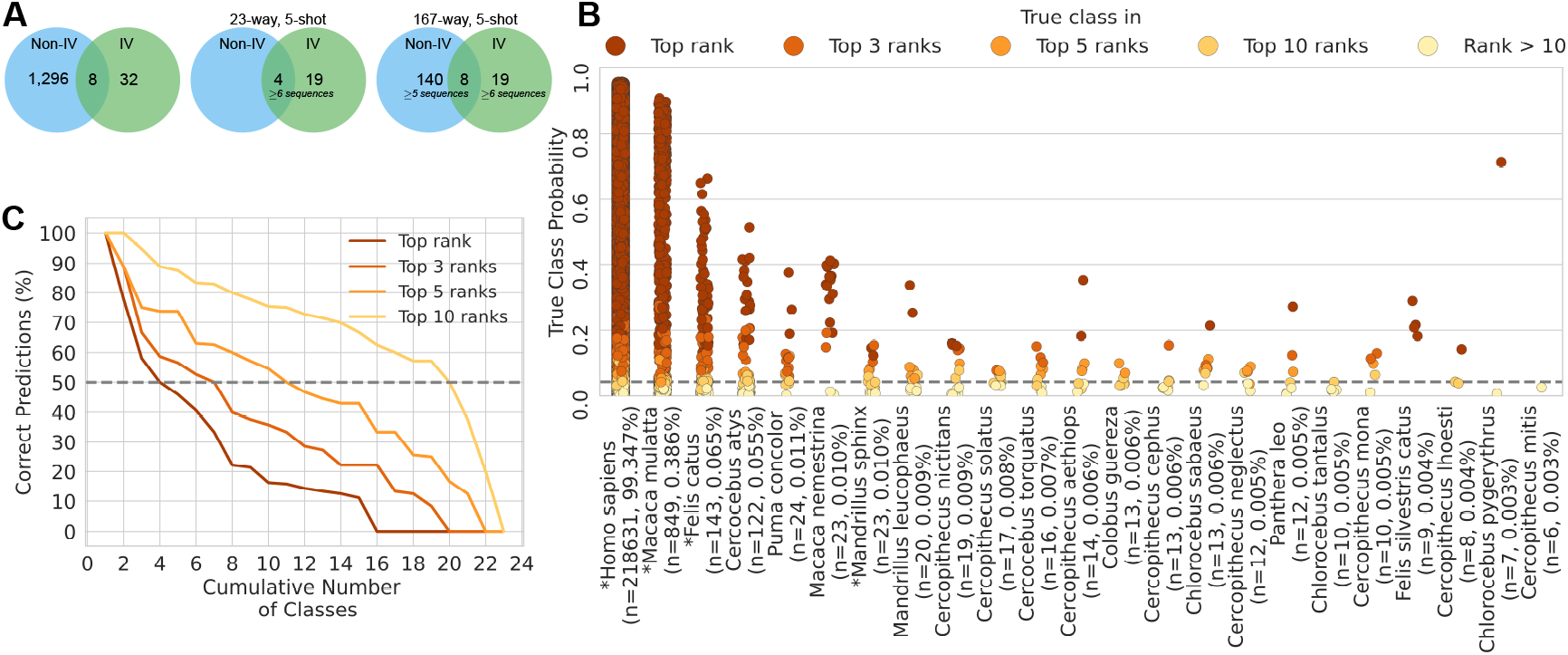
**(A)** Number of hosts from non-IV and IV datasets in 23-way and 167-way evaluations. **(B)** Distribution of the probabilities and predicted ranks of the true class for each of the 23 virus-hosts in the IV dataset in 23-way, 5-shot evaluation. An asterisk marks the four seen hosts. The vertical dashed line denotes the probability of a random predictor (= 1*/*23) for any class. **(C)** Percentage of the different rank categories of the correct predictions in each class versus the cumulative distribution of the number of classes. The horizontal dashed line indicates the number of classes with at least 50% of the samples predicted correctly in the different rank categories.

In four out of 23 classes, HAVEN predicted the true class at the top rank for 50.0% of the sequences (Figure 4C). When we considered the top three, the top five, and the top ten ranks, these values increased to 58.7%, 73.7%, and 88.9% of the sequences, respectively. Conversely, while only four classes had 50% of the sequences with correct predictions, this number rose to seven, eleven, and 20 when we considered the top three, the top five, and the top ten ranks, respectively. Notably, HAVEN accurately predicted the class for all samples of *Felis silvestris catus* (Cat) even though it had only nine labeled samples (0.004%) (Figure 4B).

The second setting of 167-way, 5-shot classification followed a similar approach to predicting hosts in the IV dataset. This repertoire of 167 classes consisted of 19 unseen and 148 seen hosts from the IV and non-IV datasets respectively, of which eight were common to both (Figure 4A). Supplementary Figure 7 indicates the predicted probabilities for the true class and the corresponding rank-based categories in the 167-way, 5-shot classification.

### 2.4 Case Study on *Coronaviridae* Spike proteins and SARS-CoV-2

Sections 2.1, 2.2, and 2.3 demonstrated the ability of HAVEN to predict common hosts for any virus, rare and unseen hosts, and hosts of unseen virus, respectively. We now turn our attention to assessing the ability of our model in a focused setting in the context of SARS-CoV-2. First, we retrained our model to predict the host when we gave Spike protein sequences from *Coronaviridae* as input and compared its performance to other pLMs. We then tested our model and pLMs against SARS-CoV-2 variants of concern, specifically asking if each model could correctly predict *Homo sapiens* as the host for each variant.

We constructed a dataset of all spike protein sequences of viruses belonging to the *Coronaviridae* family (Taxonomy id 11118) from UniRef90 [20]. Supplementary Note 2 describes the steps involved in annotating the viral sequences with their corresponding virus-hosts and the preprocessing pipeline. There were 814 spike protein sequences from 103 *Coronaviridae* viruses (Figure 5A) infecting 98 vertebrate hosts (Figure 5B). Similar to the common classes in the non-IV dataset (Section 3.1), we selected sequences of hosts with at least one percent prevalence to fine-tune HAVEN.

**Figure 5:**
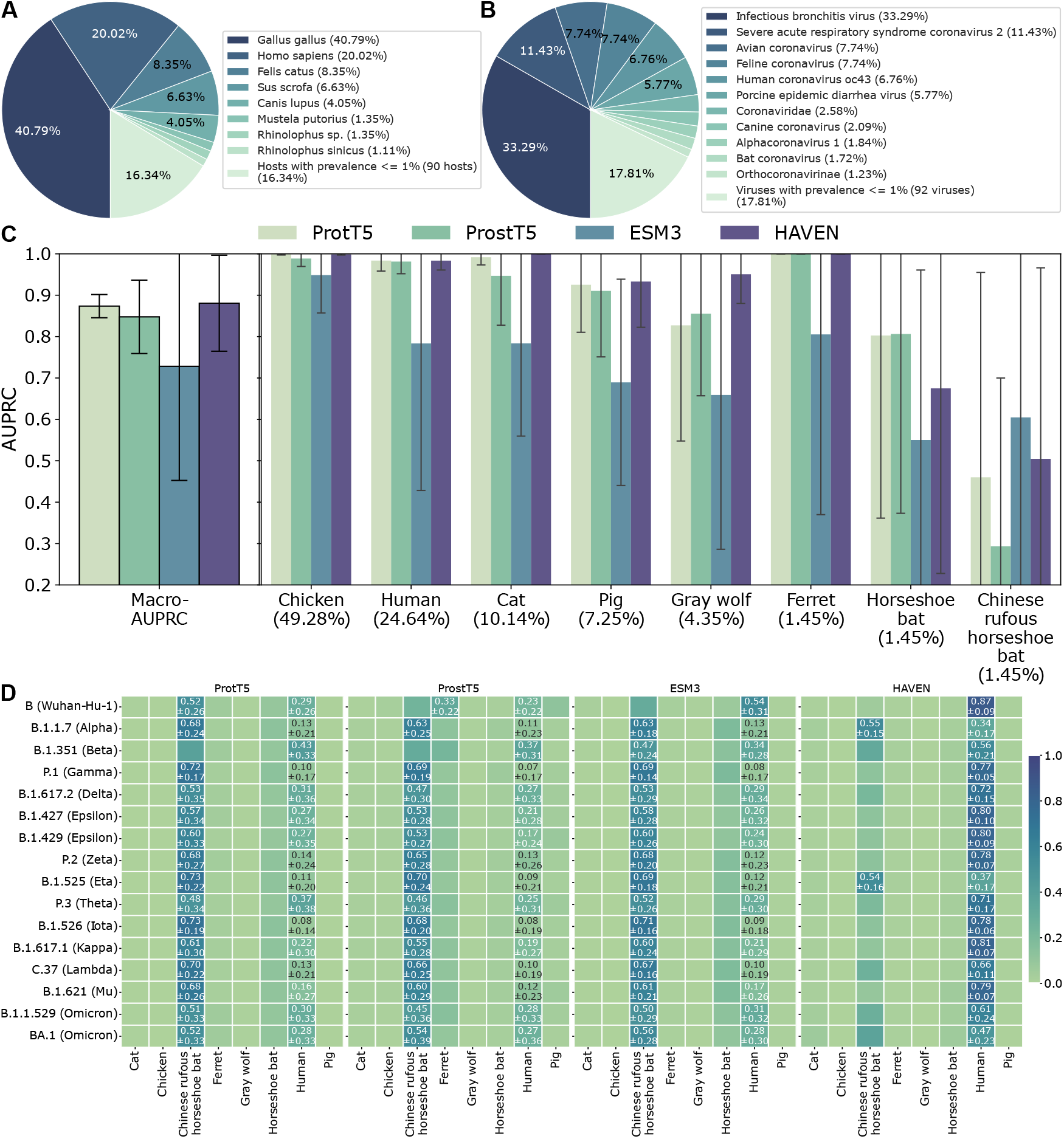
Distribution of **(A)** virus-hosts and **(B)** viruses in the dataset of Uniref90 *Coronaviridae* spike protein sequences. **(C)** Distribution of the macro-Area Under Precision-Recall Curve (AUPRC) and classwise-AUPRC scores of all models in predicting hosts for the spike protein sequences across five iterations. The height of the bars denote the mean macro-AUPRC and the error bars represent the standard deviation. **(D)** Host prediction probabilities for spike protein sequences of SARS-CoV-2 variants of concern from ProstT5, ESM2, ESM3, and HAVEN fine-tuned using Uniref90 *Coronaviridae* Spike protein datasets.

The final dataset had 681 sequences of 37 viruses sampled from eight common hosts. Starting from the version of HAVEN pre-trained on all UniRef90 viral protein sequences, we fine-tuned it using this dataset of *Coronaviridae* spike protein sequences to predict the eight classes. Similarly, we fine-tuned the baseline foundation models including ProtT5, ProstT5, and ESM3. We compared the performance of our model with these pLMs (Figure 5C). Though the differences were not significant, HAVEN had a higher mean macro-AUPRC 0.88 than ProtT5 (mean macro-AUPRC=0.87; *p*-value=1; Mann-Whitney U test), ProstT5 (0.85; 0.69), and ESM3 (0.73; 0.31).

We used this fine-tuned HAVEN and baseline pLM models to predict the hosts for 100 spike protein sequences randomly sampled from National Center for Biotechnology Information (NCBI) Virus for each of the 16 SARS-CoV-2 variants of concern (VOCs) [21, 22]. HAVEN outperformed the rest in accurately predicting the host as *Homo sapiens* (Humans) for fourteen VOCs (Figure 5D).

### 2.5 Ablation Study

We performed an ablation study to determine the importance of each of the three main components of HAVEN: pre-training, segmentation, and hierarchical self-attention. We evaluated five combinations (Supplementary Note 7): (i) without pre-training, segmentation, and hierarchical self-attention (w/o Pre-Tr, w/o Seg, w/o HSA), (ii) without pre-training and hierarchical self-attention (w/o Pre-Tr, w/o HSA), (iii) without pre-training (w/o Pre-Tr), (iv) without segmentation and hierarchical self-attention (w/o Seg, w/o HSA), and (v) without hierarchical self-attention (w/o HSA). We evaluated these models on prediction of common classes in the non-IV dataset (Supplementary Figure 3). HAVEN with mean macro-AUPRC 0.68 significantly outperformed the variants.

## 3 Methodology

### 3.1 Datasets

We describe our dataset briefly (Figure 1B, C, and D), with details in Supplementary Note 2. Our pretraining dataset contained 1, 207, 317 protein sequences, restricted to *Viridae* [20, 23]. Each sequence is guaranteed to have at most 90% similarity to any other sequence. To train the multi-class model, we obtained the host corresponding to each viral protein from European Nucleotide Archive (ENA) maintained by the European Molecular Biology Laboratory’s European Bioinformatics Institute (EMBL-EBI) [24]. Adhering to the W.H.O.’s definition of zoonoses, we retained only viruses with hosts belonging to the *Vertebrata* clade [25]. These 267, 860 protein sequences came from 3, 779 unique viruses infecting 1, 314 unique hosts. Human hosts (247, 415; 92.37%) and Human immunodeficiency virus (218, 298; 84.80%) dominated the data. To obtain a comprehensive and sufficient representation from a diverse set of classes to train HAVEN and to test its generalizability, we divided the dataset as follows:

1. **Non-Immunodeficiency Virus (non-IV) dataset**: 47, 792 sequences from the 3, 772 virus species that are not immunodeficiency viruses. These sequences are from 1, 304 hosts. Since 1, 299 hosts had less than 1% prevalence, we further partitioned this dataset into two groups: (a) *Common classes*: hosts with at least 1% prevalence. There were 31, 718 viral protein sequences from 661 unique viruses infecting five different hosts namely, *Homo sapiens* (Human), *Sus scrofa* (Pig), *Hydrochoerus hydrochaeris* (Capybara), *Marmota himalayana* (Himalayan marmot), and *Gallus gallus* (Red junglefowl). (b) *Rare classes*: hosts with prevalence at least 0.05% and less than 1%. We did not consider classes with fewer than six samples (prevalence=0.05%) to meet FSL requirements. This dataset contained 143 hosts.
2. **Immunodeficiency Virus (IV) dataset**: 220, 068 sequences of seven immunodeficiency viruses infecting 40 hosts.

We fine-tuned, validated, and evaluated HAVEN and other baseline models for multi-class virus-host prediction using the common classes in the non-IV dataset (Section 3.3). We integrated HAVEN into an FSL framework (Section 3.4) to predict rare, unseen hosts in the non-IV dataset. Finally, we used the IV dataset to assess the predictive capability on unseen viruses and with both seen and unseen hosts.

### 3.2 HAVEN Model Architecture

At its core, HAVEN is a pLM based on the BERT architecture comprising of Transformer encoder. All transformer-based language models rely on self-attention. The performance of transformer-based language models degrades for long contexts, e.g., by paying higher attention to the head and tail portions of “ long” input sequences compared to their middle parts [26]. The threshold to quantify whether a sequence is ‘long’ is subjective to the downstream task being solved and the properties of the input. Most pLMs circumvent this issue by using fixed context lengths such as 1024, 2048, or 4096 [27, 28].

The length of protein sequences in the pretraining dataset ranged from eleven to 13, 556. To avoid using a fixed context length, which would entail curtailing long sequences or discarding them from the input dataset, we included a hierarchical design into HAVEN and augmented it with self-attention [19].

The input to HAVEN was a viral protein represented as a sequence of amino acids (tokens). We encoded each amino acid token as an integer using a vocabulary of size 26. This vocabulary included 20 standard amino acids, selenocysteine (U), pyrrolysine (O), aspartic acid or asparagine (B), glutamic acid or glutamine (Z), leucine or isoleucine (J), and unknown amino-acid (X) [29, 30]. Further, there were three special tokens to represent padding, class, and mask tokens, respectively. In addition to the amino-acid token representation, we encoded the positional information of every token using sine and cosine functions [31]. The input embedding for every token in the protein sequence was a sum of the token embedding and the positional embedding.

There are two components in HAVEN (Figure 1A):

#### Segment encoder

We divided every input sequence into overlapping subsequences of fixed length (“ segments”) starting at regular intervals (“ strides”). Denoting the length of a segment by *sl* and the stride by *st*, a sequence of length *n* contained ⌊ (*n* − *sl*)*/st*⌋ +1 segments. We encoded each segment using a BERT with six identical layers of the Transformer encoder stack [31]. Each encoder layer included multi-head self-attention, a position-wise fully-connected feed forward network, and normalization layers. We prefixed every segment with a class (*CLS*) token [18]. The embedding of this *CLS* token served as the representation vector for every segment.

#### Sequence encoder

For every input sequence, the input to this component included the representation of the constituting segments from the segment encoder. HAVEN transformed these segment embeddings using a second multi-head self-attention layer thereby enabling every segment to draw information from all other segments in the sequence. Finally, we averaged the transformed embeddings of all segments to compute the representation of the entire protein sequence.

### 3.3 Pre-training and Fine-tuning HAVEN to Predict Common Classes

#### Pre-training

We pre-trained the segment encoder using the self-supervised task called masked language modeling [18]. We used a segment length of 256 with padding, if necessary, and stride of 64. For each segment, we selected 15% of the positions uniformly at random for masking. We excluded positions with *CLS* and padding tokens from masking. For each selected position, we replaced the corresponding token with (i) an explicit *MASK* token with 80% probability, (ii) a random token with 10% probability, (iii) left it unchanged with 10% probability. We pre-trained to optimize the cross-entropy loss in predicting the original token in the input sequence at the masked positions. We used 90% of the dataset for training and the remaining 10% for validation. See Supplementary Note 4 for more details.

#### Fine-tuning

We fine-tuned the pre-trained segment encoder for multi-host prediction of common classes in the non-IV dataset (Section 3.1). The sequence encoder generated embeddings for the entire protein sequence which we further processed through linear layers to compute probabilities for multiple classes. We optimized cross-entropy based focal loss [32] to account for class imbalance (Supplementary Note 5).

#### Evaluation

We compared the performance of HAVEN on predicting common classes in the non-IV dataset to vision-based, natural language processing (NLP), and foundation pLM models (Supplementary Note 6). During fine-tuning, we trained the models using 80% of the dataset, validated with another 10% to determine early-stopping, and evaluated using the final 10% as the test dataset. We measured the area under precision-recall curve (AUPRC) for each class and the macro-AUPRC (average of the per-class AUPRC). Other metrics such as accuracy, precision, recall, area under receiver operating characteristic are susceptible to over-estimating the performance of models predicting correctly for the majority class. For every fine-tuning experiment, we performed five runs using different splits of the dataset.

### 3.4 Generalizing HAVEN to Rare, Unseen Hosts and Unseen Viruses

We integrated HAVEN into a few-shot learning (FSL) technique called ‘Prototypical Networks’ [33] to enable predictions for rare and unseen hosts, and unseen viruses. In this framework, we built an *N*-way-*K*-shot FSL classifier, where *N* represents the number of classes and *K* is the number of labeled, “ support” samples in each class. The classifier uses another ML model (e.g., HAVEN) to compute an embedding for each support sample. It then computes a prototype embedding for each class as the mean of the embeddings of the support samples. For every unlabeled “ query” sample, the classifier computes a probability for the *N* classes using the Euclidean distance between this sample’s embedding and the prototypes of each class. The probability for a class is inversely proportional to the distance between the sample and the corresponding prototype. In our approach, the FSL classifier used the output of HAVEN’s sequence encoder (Figure 1A) to compute embeddings of support and query sequences. This integration enabled two types of generalizations:

#### Predicting rare and unseen hosts in the non-IV dataset

The five common classes seen by HAVEN during training were not sufficient to capture the diversity of the 143 rare, unseen hosts. Hence, we used the rare classes dataset to further fine-tune HAVEN using FSL (Supplementary Note 8). We trained and evaluated FSL classifiers with varying values of *N* and *K* (Supplementary Figures 4, 5, and 6) and selected the best performing 3-way, 5-shot model for the next generalization analysis.

#### Predicting hosts of unseen viruses in the IV dataset

Here, we used the integration of HAVEN within the FSL framework in a purely evaluative setting, i.e., we did not train or fine-tune any component of HAVEN further with any of the sequences in the IV dataset. We designed two FSL experiments using the IV dataset to mimic different real-world circumstances (Figure 4A):

1. 23**-way**, 5**-shot**: At the onset of an endemic, only a limited number of labeled samples are available for a novel virus. However, the family to which a virus belongs may enable us to focus on a specific group of hosts. Hence, we restricted our attention to animals that immunodeficiency viruses are known to infect. Our data contained 23 such hosts that also had at least six samples, permitting FSL analysis. Therefore, we created a 23-way, 5-shot classifier to predict hosts of unseen immunodeficiency viruses.
2. **167-way, 5-shot**: In a more general scenario, we may not be able to narrow our focus to a small set of hosts. Hence, we expanded the previous analysis by adding 148 hosts from the non-IV dataset that had at least five sequences; of these four hosts were also present in the IV dataset, yielding a total of 167 hosts. The generality of this framework permits the addition of a new host (with at least five viral sequences) to the model without any explicit fine-tuning.

In both analyses, the query set included only sequences from the IV dataset. Additional details of these analyses appear in Supplementary Notes 8 and 9.

## 4 Discussion

The pace of discovery of new animal viruses has accelerated in recent years, especially through metagenomic profiling of clinical and environmental samples [34, 35, 36, 37]. However, we do not know the host range for many of these viruses. Experimentally determining the host range for these viruses is time-consuming and expensive. Computational predictive models that can accurately and robustly predict the organisms infected by a virus have the potential to fill this gap. These observations motivated us to develop HAVEN and to evaluate its generalizability to unseen hosts and viruses.

Two salient and novel factors empowered HAVEN in virus-host prediction. First, the model architecture comprising segmenting the input sequence and hierarchical self-attention facilitated HAVEN to learn superior embeddings for protein sequences of any length. Second, the comprehensive pre-training dataset of viral protein sequences was highly effective. In contrast, viruses are insufficiently represented in the protein sequence databases used to train pLMs [38, 39]. These two factors optimized the number of parameters (18M) and the size of the pretraining dataset (1.2M) for HAVEN. pLMs have sizes and requirements that are several magnitudes larger, e.g., ProtT5 (1.2B parameters, 45M sequences), ProstT5 (1.2B, 35M), ESM3 (98B, 3.15M). Nevertheless, HAVEN achieved prediction performance for common classes that was on par with pLMs.

In a case study of SARS-CoV-2, HAVEN surpassed foundation pLMs in predicting hosts of variants of concern (VOCs) (Figure 5). ProtT5, ProstT5, and ESM3 incorrectly predicted the highest probability for Chinese rufous horseshoe bat for 14 out of 16 VOCs, whereas our model was inaccurate for only two VOCs. SARS-CoV-2 has ancestral origins in this bat [40]. pLM-based predictors were unable to differentiate between similar elements in the spike proteins of coronaviruses infecting this bat species and those of VOCs. Interestingly, the two VOCs for which our model performed poorly – Eta (B.1.525) and Alpha (B.1.1.7) have mutations within or near the spike protein furin cleavage site, a domain in which mutations have been associated with host range in labeled coronavirus datasets [41, 42].

We demonstrated that HAVEN can generalize to rare, unseen hosts and unseen viruses (Sections 2.2, 2.3). A potential direction for improving sequence-based prediction methods is to couple them with network-based methods [9, 11, 13, 14] or to leverage structural information on viral proteins.

## 5 Acknowledgments

National Science Foundation awards CCF-2412389, CCF-2200045, the Pandemic Prediction and Prevention Destination Area at Virginia Tech, and the College of Engineering at Virginia Tech supported this research work.

We are grateful to Jassi Pannu and Moritz Hanke from the John Hopkins Center for Health Security for reviewing this study and providing input on biosafety/biosecurity issues.

## 6 Code Availability

The software is available for public use under the GNU General Public License v3 at https://github.com/NSF-COMPASS-Center/HAVEN.

## 7 Data Availability

All the datasets used can be downloaded from https://doi.org/10.5281/zenodo.15540220. The pre-trained and fine-tuned models are accessible at https://doi.org/10.5281/zenodo.15537800. Both resources are available under the Creative Commons Attribution 4.0 International license.

