## Supplementary Material for "HAVEN: Hierarchical Attention for Viral protEin-based host iNference"

#### Supplementary Notes

#### 1 Related Work

Iuchi et al. [1] provided an extensive review of computational methods for predicting hosts of animal and avian viruses and bacteriophages. The methods ranged from standard machine learning (ML) classifiers such as k-nearest neighbors, support vector machines, and random forests to natural language processing (NLP) models [2, 3, 4]. The inputs to the feature-based ML methods included nucleotide composition, amino acid (AA) composition, motif frequency, distribution of AA groups, conjoint triads, physiochemical properties of AAs, and k-mer profiles of the viral sequences. Existing studies can be broadly categorized into three types namely, (i) binary classification, (ii) multi-label classification, and (iii) link prediction in a network.

A majority of the studies treat virus-host prediction as a binary classification task. DeePaC is an ensemble of Siamese-based networks comprising convolution and bidirectional long short-term networks [5]. This model learned information from bacterial genome sequences and their reverse complements to predict the potential of bacteria to infect humans. The authors extended the study to predict the host of a virus as human or non-human using DeePaC-vir [6]. Mollentze et al. [7] implemented gradient boosted machine learning classifiers to assess the risk of animal viruses infecting humans. They trained the classifier using viral genomic information such as codon usage biases, amino acid biases, and dinucleotide biases, and the similarity between viral and human genes. Tseng et al. [8] used boosted regression trees to classify different mammals as hosts of orthopoxovirus species. They leveraged viral genomic features and host ecological traits. VirusBERTHP is another binary classifier predicting the potential of a virus to infect humans. Each token in this NLP model corresponded to a $k$-mer in the viral genomic sequence [9]. In EvoMIL, Liu et al. [10] conceived each virus as a bag of protein sequences. They obtained embeddings for each sequence using a protein language model (pLM). For each host of interest, the authors trained a new attention-based multiple-instance learning classifier. Each of these binary classifiers predicted the association between a virus bag and a given host.

A small number of recent models have the capability to predict multiple host-labels, specifically spanning different taxonomic ranks for a given virus [11, 12]. Taking a viral genome sequence as input, RNAVirHost [11] created a two-layer XGBoost model to predict the host across different taxonomic ranks. Each layer solved a multiclass classification problem. The first layer predicted the host at the kingdom and phylum level, while the second layer focused on the class and order levels. This model used two types of features: genomic traits and related virus groups identified based on sequence alignment. The authors trained independent

RNAVirHost models for different orders of viruses. VirHRanger [12] is an ensemble of six hierarchical classifiers, each exploiting different types of information about viruses, such as genome and protein sequences, sequence composition, and human-virus protein-protein interaction networks. For a given input, each of the six hierarchical classifiers predicted host labels that spanned different taxonomic ranks from phylum to species. The ensemble architecture aggregated the predictions from the individual classifiers using an independent linear regressor for each label.

The third category of studies framed virus-host prediction as a link prediction problem in a virus-host network [4, 13, 14, 15]. Wardeh et al. [13] developed a framework with three approaches to predict new links between known viruses and mammalian hosts. The first method adopted a local, mammalian perspective: the authors trained independent binary classifiers for each mammalian species to predict associations between it and a set of known viruses, using binary, categorical, and continuous features describing viruses. The second method took a local, viral perspective using binary classifiers for every virus. The final global approach used motifs from the virus-host network to predict missing links. In each approach and for every species, they selected the best performing model from eight support vector machine, neural network-based, and tree-based models. They aggregated the findings from all three approaches to make final predictions on novel virus-mammal links. Another study [14] also predicted missing edges in a virus-mammal network using a feature-agnostic method called LF-SVD. This model combined linear filtering and singular value decomposition and leveraged network structure parameters such as in-degree, out-degree, and connectivity to impute the virus-host network. Pandit et al. [15] used network-based metrics as node features to train binary and multi-class, multi-label gradient boosting classifiers to predict links (labeled with hosts) between novel viruses and known viruses. VHIP [4] predicted virus-host interactions in a manually curated Virus Host Range network (VHRnet). This network contained virus-host pairs and their corresponding sequences. The authors trained a gradient boosting model using homology and composition features from sequences of virus-host pairs that captured information about putative hosts and coevolution signals. The features included motifs associated with CRISPR activity and horizontal gene transfer between virus and hosts, $k$-mer profile similarities, and their GC content.

Existing approaches have three key limitations. First, most of these models were trained on data from a single viral species or family. Thus, they lack the capacity to generalize to previously unseen viruses or hosts. Second, many methods are restricted to binary classification tasks: determining whether a virus can infect a specific host (often humans) or not. Hence, these approaches are not applicable to predicting the animal or reservoir host of a virus in nature. Finally, the multi-label and network-based approaches [4, 11, 12, 13, 14] were trained on virus-host networks obtained from sources such as the Virus-Host Database [16] and VIRION [17]. An edge in these networks indicates a association between a virus and a host at the organism level. Therefore, these models cannot predict the host given an individual protein sequence of a virus.

These observations motivated us to develop a virus-host predictor that addresses all these research gaps. The model should be trained on a comprehensive dataset of sequences spanning a broad range of viruses and host organisms. Given a viral protein sequence, the model should solve a multi-class classification task. This model should also generalize to make accurate predictions for unseen hosts and viruses not seen during the training process.

#### 2 Dataset Sources and Construction

We constructed our dataset in four steps: (i) downloading a comprehensive and diverse set of viral protein sequences, (ii) curating a dataset of these sequences for pretraining HAVEN, (iii) obtaining host labels for each sequence, and (iv) defining different subsets for fine-tuning HAVEN to predict common and rare hosts of seen and unseen viruses. Supplementary Table 1 provides statistics on our data after each of these steps.

**Viral protein sequences.** An important consideration in our choice of dataset was that the presence of highly similar protein sequences in both the training and test datasets may lead to data leakage and over-estimation of model performance. Hence, we considered UniProt Reference Clusters (UniRef) in UniProt Knowledgebase (UniProtKB) [18]. These are clusters of protein sequences grouped together based on sequence identity. There is a representative sequence for each cluster [19]. UniRef100 combines identical sequences whereas UniRef90 combines UniRef100 seed sequences with at least 90% sequence identity and 80% overlap with the longest sequence in the cluster. The number of representative proteins in UniRef90 is 42% of the number of proteins in UniProt [20]. We downloaded these representative protein sequences of UniRef90 clusters for our analysis. We refer to them as UniRef90 sequences henceforth.

**Dataset for pre-training.** As of January 31, 2024, there were $184,520,054$ UniRef90 sequences. We used the UniProtKB web interface to select the reference sequences of $1,257,643$ UniRef90 clusters belonging to the *Viridae* family, corresponding to the National Center for Biotechnology Information (NCBI) taxonomy ID 10239 [21]. There were $1,207,317$ sequences after removing the ones that did not have a valid taxonomy identifier for the virus. We pretrained HAVEN using this dataset of viral UniRef90 sequences.

**Host labels of protein sequences.** The "virus hosts" field in every UniProtKB [18] protein record indicated either a specific organism or taxonomic group of organisms that are susceptible to be infected by the virus. This value does not necessarily denote the specific organism from which the viral genome sequence was originally sampled. The "protein_id" present in the metadata of each protein record in UniProtKB identifies the source sequence in the European Nucleotide Archive (ENA) maintained by the European Molecular Biology Laboratory's European Bioinformatics Institute (EMBL-EBI). We used EMBL-EBI's 'database fetch (Dbfetch)' tool to retrieve the "host" field in the record for this source sequence [22, 23]. If this value was not empty, we converted it into an NCBI taxonomy identifier using TaxonKit [24]. We used this identifier as the host label of the protein sequence.

**Datasets for fine tuning.** We cleaned the dataset of $1,257,643$ viral UniRef90 sequences by removing sequences based on the following criteria:

1. there was no NCBI taxonomy identifier associated with this sequence (this situation arose for about $50,000$ sequences even though we had used *Viridae* as the selection criterion in the UniProtKB web interface),
2. there was no EMBL-EBI reference identifier,
3. the sequence mapped to the EMBL-EBI reference for another sequence (duplicates),
4. the "host" field in ENA was empty,
5. the "host" field was not empty but TaxonKit was not able to map it to a valid NCBI taxonomy identifier. Examples of such host values are "Castanea sativa x Castanea crenata) x Castanea mollissima cv. Precoce Migoule CA127", "Homo sapiens; sex: M; age: 7 months", and "Streptococcus pasteurianus DA586".

We applied two filtering criteria to curate the fine-tuning dataset. We retained a sequence in the cleaned dataset only if -

1. both the virus and host organisms were specified at the "species" NCBI taxonomy rank,
2. the host species belonged to the *Vertebrata* clade [25].

The resulting dataset consisted of $267,860$ protein sequences from $3,779$ unique viruses infecting $1,314$ unique hosts. Notably, this dataset was predominantly composed of sequences from human hosts (92.37%) and Human immunodeficiency viruses (84.80%). We further divided the dataset to fine-tune HAVEN using a diverse set of classes and evaluate its generalizability to rare and unseen hosts, and unseen viruses:

1. Non-immunodeficiency virus (non-IV) dataset ($47,792$; 21.72%): sequences from virus species that were not immunodeficiency virus species. This dataset included $3,772$ virus species infecting $1,304$ hosts (Supplementary Figure 1a).
   (a) Common hosts dataset: $31,718$ viral protein sequences from 661 unique viruses infecting five virus-hosts with atleast 1% prevalence in the Non-IV dataset. The hosts included *Homo sapiens* (Human), *Sus scrofa* (Pig), *Hydrochoerus hydrochaeris* (Capybara), *Marmota himalayana* (Himalayan marmot), and *Gallus gallus* (Red junglefowl). The lengths of the sequences in this dataset ranged from eleven to $7,217$ with 93.1% of the sequences shorter than 612 amino acids. To meet the limitations of the computational resources, we set a cut-off at 99.9 percentile of the sequence length. Thus, only sequences with length $\leq 3,036$ were considered for fine-tuning HAVEN.

(b) Rare hosts dataset: $16,074$ sequences of 143 hosts with $0.05\% \leq$ prevalence $< 1\%$. We selected hosts with atleast six samples to allow for few-shot learning (FSL) analysis. We further pruned this dataset to select only sequences with length within the 99 percentile which set the cut-off as sequence length $\leq 2,452$.

2. Immunodeficiency virus (IV) dataset ($220,068$; prevalence=82.16%): sequences from the seven Immunodeficiency viruses enlisted in Supplementary Table 2 (Supplementary Figure 1b).

| **Description** | **# sequences** | **# unique viruses** | **# unique virus-hosts** |
|---|---|---|---|
| UniRef90 clusters | $184,520,054$ | – | – |
| Viral protein sequences (NCBI Taxonomy ID = 10239) | $1,257,643$ | – | – |
| AND with valid virus taxonomy ID | $1,207,317$ | – | – |
| AND with valid EMBL-EBI identifier | $1,184,006$ | – | – |
| AND with valid host organism from ENA) | $613,208$ | – | – |
| AND virus taxonomy rank = species | $463,755$ | – | – |
| AND virus-host taxonomy rank = species | $348,401$ | $10,967$ | $3,662$ |
| AND virus-hosts belonging to the *Vertebrata* clade | $267,860$ | $3,779$ | $1,314$ |
| Immunodeficiency virus protein sequences | $220,068$ | 7 | 40 |
| NOT immunodeficiency virus protein sequences | $47,792$ | $3,772$ | $1,304$ |
| AND virus-hosts with at least 1% prevalence | $31,718$ | 661 | 5 |

Table 1: Number of sequences, unique viruses, and unique virus-hosts at various stages of dataset construction.

| **Virus Name** | **# Sequences** | **# Prevalence in IV dataset** |
|---|---|---|
| Human immunodeficiency virus 1 | $215,415$ | 97.89% |
| Human immunodeficiency virus | $2,095$ | 0.95% |
| Simian-Human immunodeficiency virus | $1,030$ | 0.47% |
| Human immunodeficiency virus 2 | 788 | 0.36% |
| Simian immunodeficiency virus | 538 | 0.24% |
| Feline immunodeficiency virus | 198 | 0.09% |
| Bovine immunodeficiency virus | 4 | 0.001% |

Table 2: Composition of the immunodeficiency virus (IV) dataset

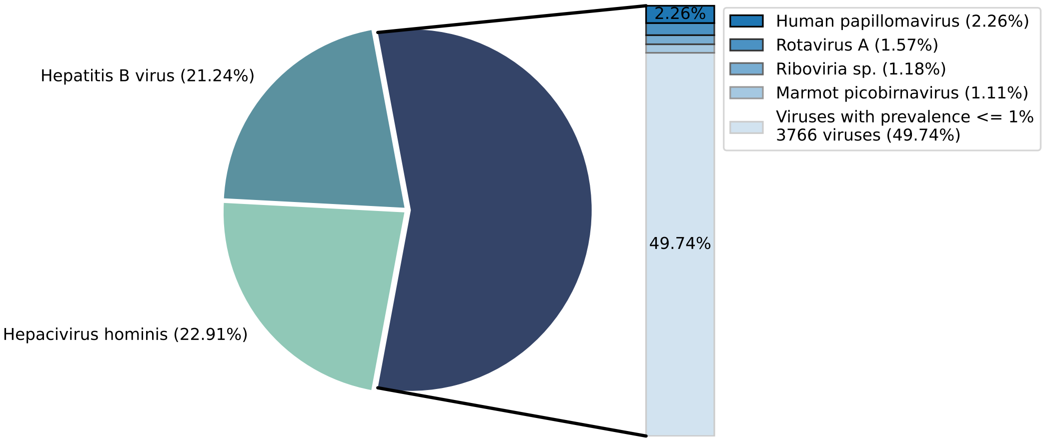


(a) Non-immunodeficiency Virus Dataset

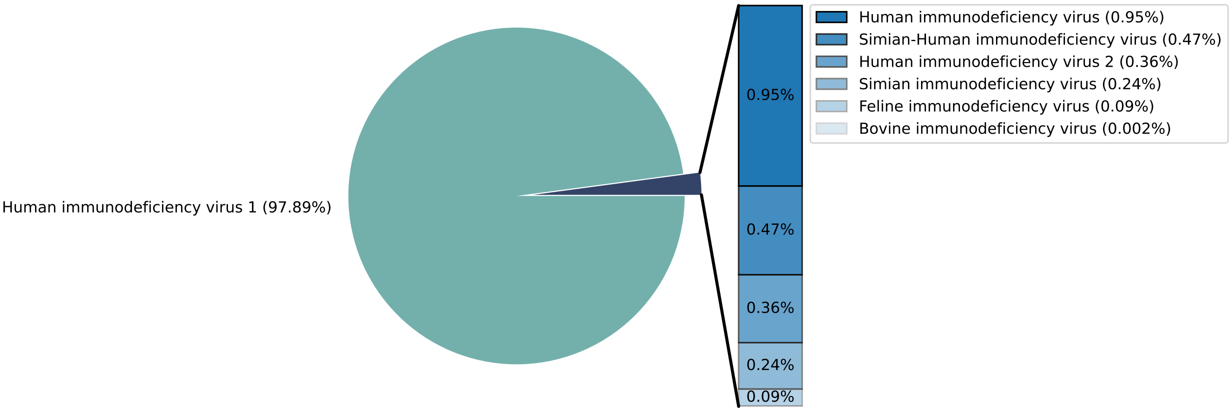


(b) Immunodeficiency Virus Dataset

Figure 1: Distribution of viruses in the (a) non-immunodeficiency Virus (Non-IV) and (b) immunodeficiency Virus (IV) Datasets. The non-IV dataset consisted of sequences from $3,772$ viruses of which $3,766$ viruses had $\leq 1\%$ prevalence in the dataset. The IV dataset comprised of seven viruses with Human immunodeficiency virus 1 (97.89%) dominating the dataset. The remaining six viruses had $< 1\%$ prevalence in the dataset.

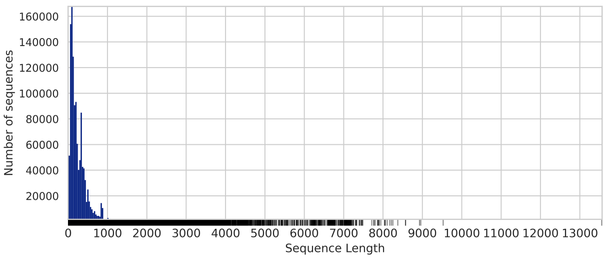


(a) Pretraining Dataset

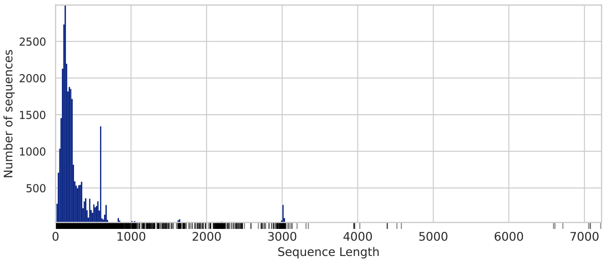


(b) Fine tuning Dataset

Figure 2: Distribution of the length of sequences in the **(a)** pretraining and **(b)** fine tuning datasets. Each tick along the $x$-axis denotes a distinct sequence length. The fine tuning dataset included viral protein sequences of the five common hosts in the non-immunodeficiency viruses (common hosts dataset).

#### 3 Hyperparameter Selection

As noted in the main text, HAVEN is a protein language model (pLM) that integrates the Bidirectional Encoder Representations from Transformers (BERT) [26] architecture with hierarchical self-attention. We used Rectified Linear Unit (ReLU) activation in all layers of HAVEN [27]. We performed an extensive hyperparameter search independently for each of the following key parameters in the model configuration. We state the parameters and the different values considered for tuning:

1. Techniques to compute positional embedding for every input token in the protein sequence: (i) index of the token in the sequence, (ii) sine and cosine functions of different frequencies [28].

2. Size of the intermediate feature vectors representing the input sequence between different layers in the model architecture: $[512, 1024]$.

3. Techniques to represent a segment in the input sequence: (i) embedding of the $CLS$ token prefixed in every segment, (ii) average of the embeddings of all amino acid tokens in the segment.

4. Maximum learning rate used to optimize the model parameters: $[1 \times 10^{-1}, 1 \times 10^{-2}, 1 \times 10^{-3}, 1 \times 10^{-4}, 1 \times 10^{-5}, 1 \times 10^{-6}, 1 \times 10^{-7}]$

5. Number of intermediate linear layers in the MLP used for multi-class classification: $[2, 4, 6, 8]$

#### 4 HAVEN Training

For both pre-training and fine-tuning, we used the Adam optimizer with a scheduled learning rate with cosine annealing [29]. This included increasing the learning rate to predefined maximum learning rate for the first 10% of the training steps followed by decreasing the learning rate to a minimum of $10^{-4}\times$ its maximum. We employed an early stopping criterion based on validation loss to prevent overfitting during pre-training and fine-tuning. Specifically, we halted the training process if the validation loss did not decrease for ten consecutive epochs and selected the model with the lowest validation loss (from all epochs) for evaluation.

**Pre-training.** We divided every input sequence into segments of length $sl = 256$ (padding, if necessary) and prefixed each segment with the $CLS$ token. We used all viral protein sequences from UniRef90 ($n = 1,207,317$) to pre-train the segment encoder using masked language modeling (MLM). We trained for 50 epochs with a batch size of 512. The maximum learning rate was $1 \times 10^{-4}$. The pre-training duration was 98 hours using one node in one HPE Apollo 6500 GPU.

**Fine-tuning.** Similar to pre-training, we trained for 50 epochs with a batch size of 32. We froze the pre-trained segment encoder for the first 20 epochs, i.e., we updated only the weights of the sequence encoder and fine-tuning linear layers during back-propagation. Subsequently, we released all the components and updated the parameters of both the segment and sequence encoders for another 30 epochs. We set the maximum learning rate to $1 \times 10^{-4}$.

**Implementation.** We used Python v3.11.9 to implement HAVEN. The machine learning and deep learning models were implemented using scikit-learn v1.5.0 and torch v2.3.1 respectively. We used seaborn v0.13.2 to create the visualizations. All pre-training and virus-host prediction experiments were performed using one node of one HPE Apollo 6500 GPU. We used one node with three HPE Apollo 6500 GPUs for FSL experiments. We pretrained HAVEN (18 million parameters) over 1.2 million viral protein sequences for $\sim$ 98 hours (4.09 days) using one HPE Apollo 6500 GPU.

#### 5 Focal Loss

As noted above, our fine tuning dataset had a high degree of class imbalance (Supplementary Note 2). The majority class could bias the learning process. We used a modified version of binary cross-entropy loss named "Focal loss (FL)"[30], which can be extended for multi-class settings, to address data imbalance and decrease the possibility of learning a model optimized to predict only the majority class correctly.

In multi-class classification, the cross-entropy loss for sample $i$ is defined as follows:

$$CE(\mathbf{y}^{(i)}, \mathbf{p}^{(i)}) = -\sum_{c \in C} y_c^{(i)} \log p_c^{(i)},$$

where $C$ is the set of classes, $p_c^{(i)}$ is the probability predicted by the model that sample $i$ belongs to class $c$, $\mathbf{p}^{(i)}$ is the vector of predicted class probabilities for sample $i$, $y_c^{(i)} = 1$ if $c$ is the true label for sample $i$ and 0 otherwise, and $\mathbf{y}^{(i)}$ is the vector of true class labels for sample $i$. Note $y_c^{(i)}$ will be zero for all classes except for the true class.

Weighted cross-entropy loss tackles class imbalance by introducing a weighting factor $\alpha_c$ for each class $c$. This weighting factor is inversely proportional to the class frequency. FL further introduces a focusing parameter $\gamma \geq 0$, which seeks to down-weight the loss for a well-classified sample and up-weight the loss for a misclassified sample. In our study we set $\gamma = 2$. The FL for for a given sample $i$ is defined as follows:

$$FL(\mathbf{y}^{(i)}, \mathbf{p}^{(i)}) = -\sum_{c \in C} \alpha_c \left(1 - p_c^{(i)}\right)^{\gamma} y_c^{(i)} \log p_c^{(i)}$$

### 6 Baseline Models

Along with HAVEN, we trained and evaluated multiple vanilla deep learning models and foundation pLMs for virus-host prediction. The vanilla models included logistic regression (LR), random forest (RF), and support vector machine (SVM), fully connected neural networks (FNN), vision-based models such as convolution neural network (CNN) adapted for textual inputs, and language-based models such as recurrent neural network (RNN), and long-short term memory (LSTM). We also fine-tuned state-of-the-art pLMs such as ProtT5 [31], ProstT5 [32], and ESM3 [33].

**Feature-based models.** We trained LR, RF, and SVM models using amino acid 3-mers as features. In every iteration of training and testing (five iterations for each model), we determined the features using a two-step process. We describe the process using the quantities from one such randomly picked iteration. First, we formed all possible distinct 3-mers of all unique amino acids in the training dataset. There were $9,431$ such combinations. In order to avoid a sparse feature matrix, we retained only the 3-mers occurring in atleast ten percent of the training dataset sequences resulting in 498 features. We represented each sequence using a vector of frequencies of these selected features. We normalized the features across all the sequences in the training dataset and testing dataset independently. For each feature-based model, we selected the optimal values for the model configuration using grid search based five-fold cross-validation. The configuration of each model and values considered while tuning the hyperparameters of the models are as follows:

1. **Logistic Regression (LR)** model optimized L1 (lasso) penalty using Stochastic Average Gradient Descent (saga) solver. Hyperparameter values for regularization term were $[0.01, 0.1, 1]$.

2. **Random Forest (RF)** model with number of decision tree classifiers selected from $[10, 100, 1000]$. Hyperparameter values for maximum depth of each tree were $[3, 5]$.

3. **Support Vector Machine (SVM)** model with Radial Basis Function Kernel. Hyperparameter values for the regularization term were $[0.01, 0.1, 1, 10]$.

**Deep learning (DL) models.** We selected the number of hidden layers for all the vanilla DL models to ensure that the number of parameters was of the same order of magnitude (12M to 14M) across these models. We performed hyperparamter tuning to select the dimension for intermediate feature vectors from $[256, 512, 1024, 2048]$ as well as the optimal maximum learning rate from $[1 \times 10^{-3}, 1 \times 10^{-4}, 1 \times 10^{-5}]$. Similar to the training of HAVEN (Supplementary Note 4), we trained each of the baseline models using a scheduled learning rate for a maximum of 50 epochs with early stopping if the validation loss did not decrease for more than ten consecutive epochs.

For all DL models, the final output layer consisted of a linear layer with output dimension equal to the number of label classes. This layer predicted the probability for each of the host class. Similar to HAVEN, we used ReLU activation in all the baseline deep learning models [27].

We computed the input embeddings for all the baseline DL models (FNN, CNN, RNN, and LSTM) using the amino-acid vocabulary embedding described for HAVEN.

1. **Fully-connected Neural Network (FNN)** model consisted of ten layers of a fully-connected feed forward neural network. The dimensions of the input layer and hidden layers were 512 and $1,024$ respectively. The maximum learning rate was $1 \times 10^{-3}$.

2. **Convolutional Neural Network (CNN)** model consisted of four convolution layers with kernel size 3 and stride 1. We used hyperparameter search to select the optimal kernel ($k$) and stride ($s$) configuration $(k, s)$ from $[(1,1),(3,1),(3,3)]$. The sizes of the input layer and hidden layers were 512 and $1,024$ respectively. The maximum learning rate was $1 \times 10^{-3}$.

3. **Recurrent Neural Network (RNN)** model consisted of six recurrent layers. The sizes of the input layer and hidden layers were 512 and $1,024$ respectively. The maximum learning rate $1 \times 10^{-5}$.

4. **Long-Short Term Memory (LSTM)** model consisted of two LSTM layers. The sizes of the input layer and hidden layers were 512 and $1,024$ respectively. The maximum learning rate was $1 \times 10^{-3}$.

**Protein Language Models (pLMs).** We implemented multi-class classifiers using three state-of-the art pretrained pLMs, namely, ProtT5, ProstT5, and ESM3 (Supplementary Table 3). We added a linear layer to transform the embeddings from the pLMs into output probabilities for each class. During fine-tuning, we updated the parameters of only this linear layer. We did not change the weights of the pLMs themselves.

1. **ProtT5** is a T5 sequence-to-sequence model pre-trained using MLM on 2.1 billion and 40 million protein sequences from Big Fantastic Database (BFD) and UniRef50, respectively [31]. We fine-tuned the 'ProtT5-XL-U50' model.

2. **ProstT5** is also a T5-based model built on top of ProtT5 that uses both sequence- and structure-based representations of proteins [32]. The encoder and decoder of T5 were initialized with the weights of ProtT5. The pre-training dataset consisted of 17 million high-quality and non-redundant protein sequences from UniProtKB and their corresponding structures from AlphaFold Protein Structure Database.

3. **ESM3** is a generative masked language model for biology. It was pretrained with three modalities of protein information: 3.15 billion protein sequences, 236 million structures, and 539 million functions. We fine-tuned the 'esm3_sm_open_v1' model for comparison [33].

| **Model Name** | **# of Params** | **Pretraining Dataset Size** |
|---|---:|---:|
| HAVEN | 18.4M | 1.2M |
| ProtT5 | 1.2B | 45M |
| ProstT5 | 1.2B | 35M |
| ESM3 | 98B | 3.15B |

Table 3: Comparison of sizes of HAVEN with state-of-the-art pLMs.

### 7 Ablation Study

HAVEN contains three main components: (i) pre-training a BERT architecture, (ii) segmentation of the input sequences into smaller segments, and (iii) a second layer of self-attention to aggregate the segment embeddings and compute the embedding of the protein sequence. We call these the pre-training, segmentation, and hierarchical self-attention modules, respectively. We performed an ablation study to ascertain the contribution of each of these components. Supplementary Figure 3 shows the performance of five different model architectures comprising varying combinations of these three components:

1. Without pre-training, segmentation, and hierarchical self-attention (w/o Pre-Tr, w/o Seg, w/o HSA): We used a vanilla transformer encoder to compute embeddings of viral protein sequences with a maximum context length of 1024.

2. Without pre-training and hierarchical self-attention (w/o Pre-Tr, w/o HSA): We divided the input sequence into segments of length 256 with stride 64. We used a vanilla transformer encoder to compute the embeddings of these segments. The representation for a sequence was the average of the embeddings of all the segments in the sequence.

3. Without pre-training (w/o Pre-Tr): We used the architecture of HAVEN in its entirety. However, we did not pre-train the segment encoder using MLM.

4. Without segmentation and hierarchical self-attention (w/o Seg, w/o HSA): We pretrained a BERT model using MLM. The maximum context length of the BERT model was 1024. We truncated input sequences longer than this threshold and computed the embeddings of sequences using the pre-trained BERT.

5. Without hierarchical self-attention (w/o HSA): We adopted the pre-trained segment encoder in HAVEN to compute representations of segments. The representation for a sequence was the average of the embeddings of all the segments in the sequence.

Note that hierarchical self-attention requires segmentation in the architecture. Hence, we did not need to consider ablation studies for the two models with hierarchical-self attention but (i) without pre-training and segmentation, and (ii) without segmentation.

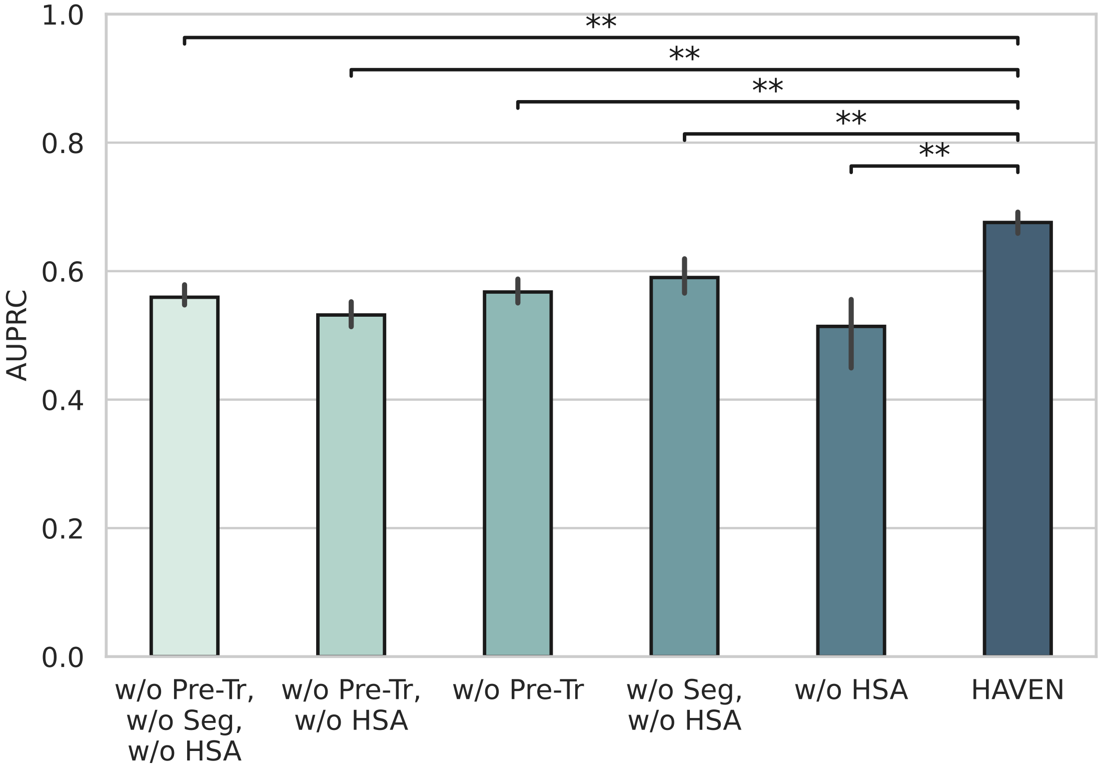


Figure 3: AUPRC scores of varying combinations of five unique combinations of different components in HAVEN architecture highlighting the contribution of each component in the model.

We evaluated the models based on their performance in host prediction in the Non-IV dataset and compared with HAVEN's performance. HAVEN with mean macro-AUPRC 0.68 significantly outperformed 'w/o Pre-Tr, w/o Seg, w/o HSA' (mean macro-AUPRC=0.56, $p$-value=$6.09 \times 10^{-3}$), 'w/o Pre-Tr, w/o HSA' (0.53,

$3.97 \times 10^{-3}$), 'w/o Pre-Tr' (0.57, $3.97 \times 10^{-3}$), 'w/o Seg, w/o HSA' (0.59, $7.94 \times 10^{-3}$), and 'w/o HSA' (0.51, $3.97 \times 10^{-3}$).

### 8 Few-Shot Learning to Predict Rare and Unseen Hosts in Non-IV dataset

We used the rare classes from the non-IV dataset containing sequences from hosts with $0.05\% \leq$ prevalence $< 1\%$ to train and test the few-shot classifier. This filter based on the class prevalence ensured that each selected class had sufficient number of samples to compose both the support and query sets for FSL. We split the dataset based on the hosts and used sequences from 70% of the rare hosts for training, 10% for validation, and the remaining 20% hosts for testing.

During the training of a $N$-way, $K$-shot 'prototypical network' classifier, every 'episode' (batch) contained $K$ support and $Q$ query sequences for each of the $N$ classes. Here $N$, $K$, and $Q$ are variables representing the number of classes, number of support sequences for each class, and number of query sequences for each class respectively. To construct every episode, we randomly sampled $N$ classes uniformly at random from the set of all rare classes and further sampled support and query sequences uniformly at random for each class.

The forward pass of a FSL classifier included three steps: (i) compute embeddings for support and query sequences in the episode using the segment encoder of HAVEN, (ii) compute the prototype for every class in the episode as the mean of the embeddings of the class's $K$ support sequences, (iii) for every query sample $x$, compute probability for each of the $N$ classes using the Euclidean distance between the embedding of $x$ and the prototype of the class. The closer the query sample is to the prototype of a class, the higher the probability for that class. During training, we used the cross-entropy loss from the predictions for the query set to update the model parameters of HAVEN through back-propagation. We repeated this training for 100 epochs where each epoch contained 100 episodes. The training process also included an early stopping criterion based on validation loss (Supplementary Note 4).

For testing the FSL we adopted a different mechanism to compose an episode. Every episode contained $N$ classes and $K$ support sequences for each class. However, unlike the training and validation phases, we selected all the remaining sequences of the selected $N$ classes as query sequences. This deviation from setting a fixed number of $Q$ query samples in each episode (unlike during training and validation) helped to retain the prevalence of the rare classes in the dataset and evaluate the model performance in an imbalanced class setting. The testing phase also constituted 100 epochs of 100 episodes each. We repeated the entire train, validate, and test pipeline five times with varying random splits of the rare classes.

Supplementary Figures 4, 5, and 6 show the performance of seven independent prototypical network classifiers trained and evaluated over the rare-class non-IV dataset with varying $N$-way, $K$-shot configurations. The mean AUPRC scores of the FSL classifiers decreased from 0.65 to 0.55 as $K$ decreased from 5 to 1 in 3-way, $K$-shot classifiers. Similarly, the mean AUPRC scores decreased from 0.65 to 0.46 as $N$ increased from 3 to 5 in $N$-way, 5-shot classifiers.

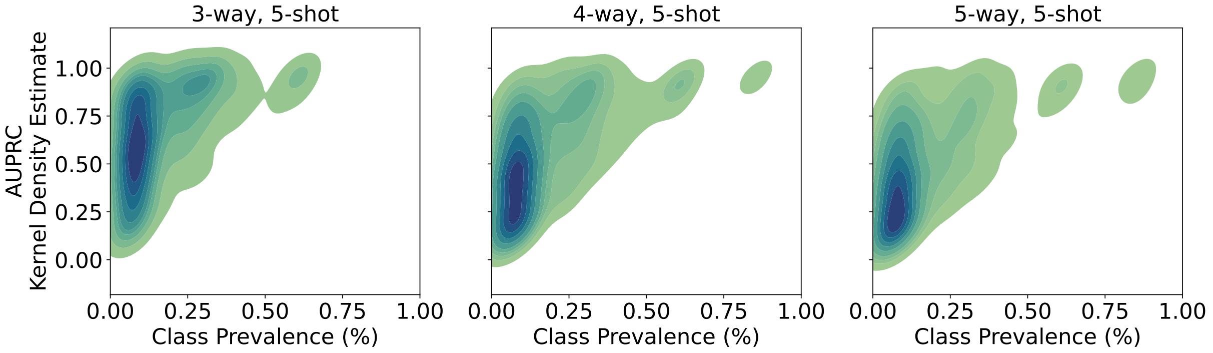


Figure 4: Bivariate kernel density estimate distribution of the AUPRC scores for each rare class against the corresponding prevalence of the class for varying number of support sequences in the $N$-way, 5-shot learning configurations. The distribution contained $50,000 \times N$ data points (five iterations, each iteration had 100 epochs, each epoch had 100 episodes, and each episode had $N$ randomly sampled rare virus-hosts). Each point denotes the AUPRC of a rare-class in one test episode ($y$-axis) and the prevalence of the class in the non-IV dataset ($x$-axis)

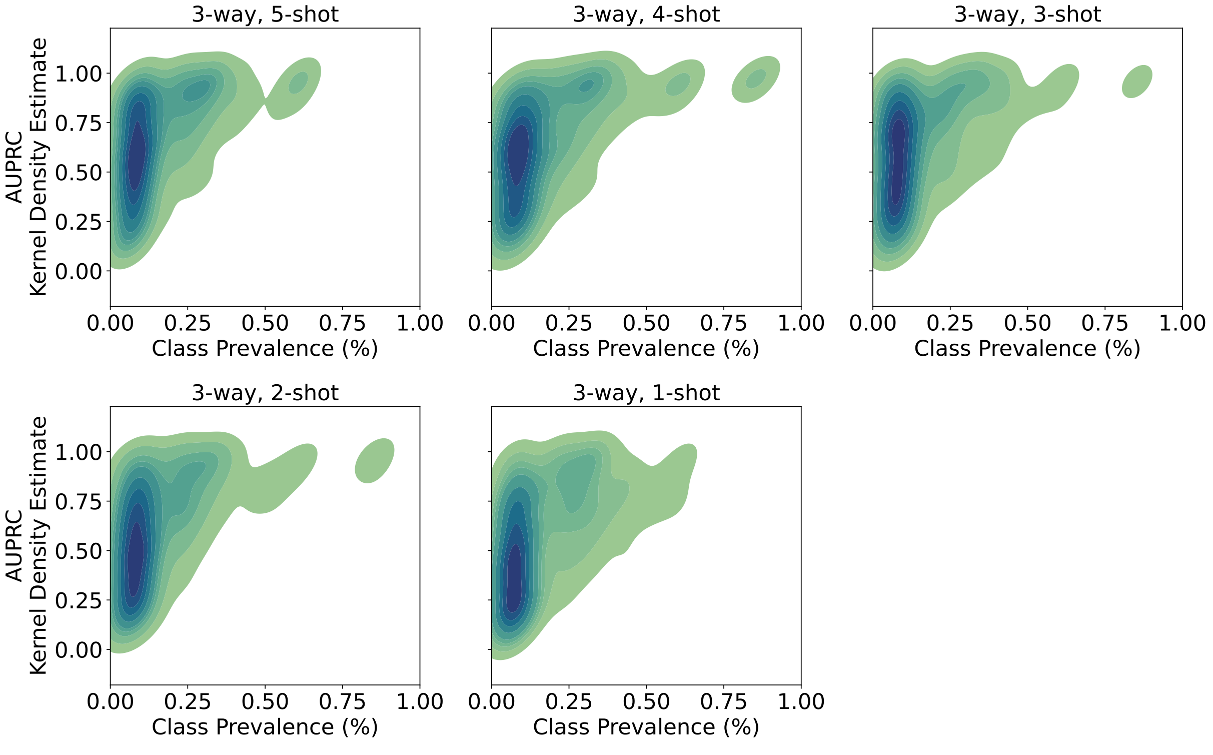


Figure 5: Bivariate kernel density estimate distribution of the AUPRC scores for each rare class against the corresponding prevalence of the class for varying number of support sequences in the 3-way, $K$-shot learning configurations. The distribution contained $150,000$ data points (five iterations, each iteration had 100 epochs, each epoch had 100 episodes, each episode had three randomly sampled rare virus-hosts). Each point denotes the AUPRC of a rare-class in one test episode ($y$-axis) and the prevalence of the class in the non-IV dataset ($x$-axis).

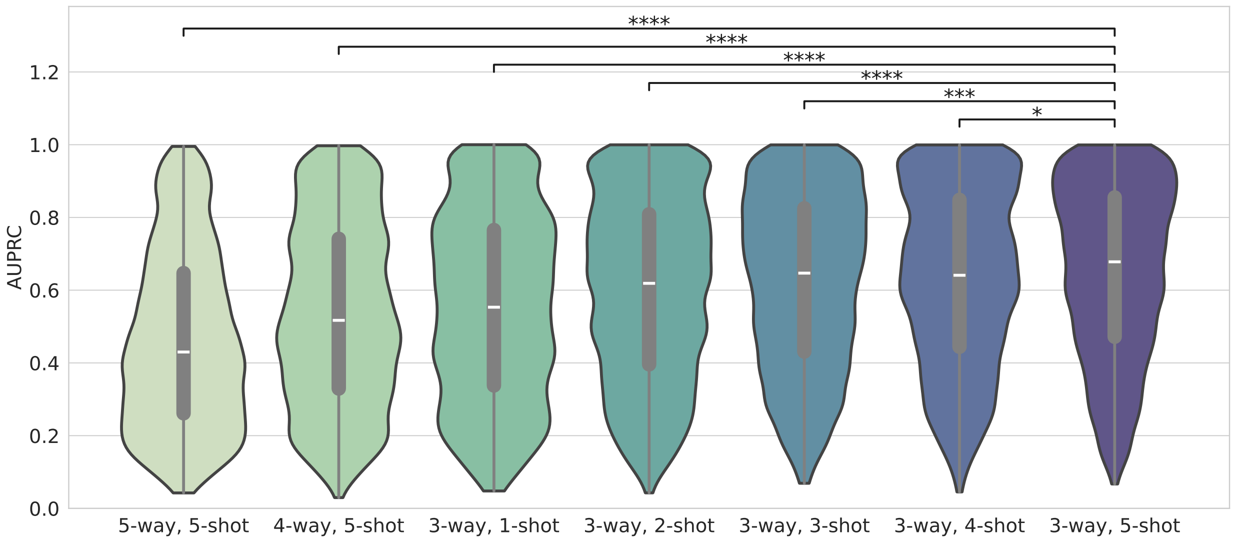


Figure 6: Distribution of AUPRC scores of all varying configurations of $N$-way, $K$-shot few-shot learning classifiers. For each model, it is a distribution of $150,000$ AUPRC scores of rare classes (five iterations, each iteration had 100 epochs, each epoch had 100 episodes, each episode had three randomly sampled rare virus-hosts). The 3-way, 5-shot classifier performed significantly better than all other combinations.

The 3-way, 5-shot (mean AUPRC=0.65) model performed significantly better than 3-way, 4-shot(mean AUPRC=0.63; $p$-value=$1.27 \times 10^{-2}$; Mann-Whitney U test), 3-way, 3-shot (0.62; $1.92 \times 10^{-4}$), 3-way, 2-shot (0.60; $9.55 \times 10^{-9}$), 3-way, 1-shot (0.55; $1.77 \times 10^{-26}$), 4-way, 5-shot (0.53; $5.25 \times 10^{-42}$), 4-way, 5-shot (0.46; $1.05 \times 10^{-108}$)
We chose the best performing 3-way, 5-shot classifier for all FSL analyses. We fine-tuned HAVEN using this FSL configuration to predict the rare and unseen hosts in the non-IV dataset.

### 9 Predicting Hosts of Unseen Virus in IV dataset

We evaluated the ability of HAVEN to predict hosts of unseen viruses using the FSL classifier fine-tuned on rare classes in the non-IV dataset. We designed a 167-way, 5-shot experiment in which the FSL classifier computed a probability for 167 hosts (148 seen and 19 unseen) for every sequence in unseen IV dataset (Supplementary Figure 7). The support set for the seen and unseen hosts included five random sequences from the non-IV and IV datasets respectively. The only requirement for adding a new host to this repertoire of 167 hosts is that there are at least five viral sequences (irrespective of which virus) known to infect the given new host.

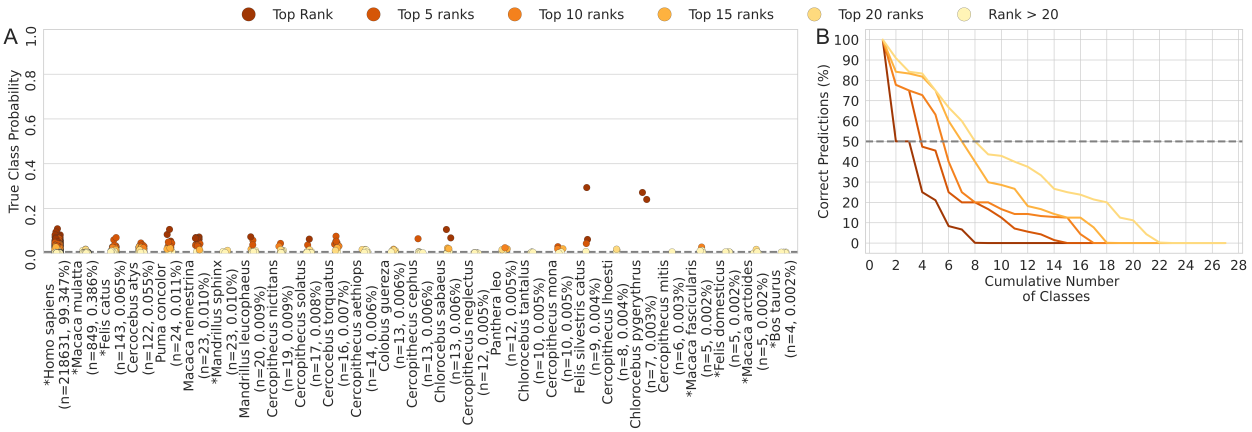


Figure 7: **(A)** Distribution of the probabilities and predicted ranks of the true class for each of the 167 virus-hosts in the IV dataset in 167-way, 5-shot evaluation. An asterisk marks the four seen hosts. The vertical dashed line denotes the probability of a random predictor (= 1/167) for any class. **(B)** Percentage of the different rank categories of the correct predictions in each class versus the cumulative distribution of the number of classes. The horizontal dashed line indicates the number of classes with at least 50% of the samples predicted correctly in the different rank categories.

[15] Pranav S. Pandit, Simon J. Anthony, Tracey Goldstein, Kevin J. Olival, Megan M. Doyle, Nicole R. Gardner, Brian Bird, Woutrina Smith, David Wolking, Kirsten Gilardi, Corina Monagin, Terra Kelly, Marcela M. Uhart, Jonathan H. Epstein, Catherine Machalaba, Melinda K. Rostal, Patrick Dawson, Emily Hagan, Ava Sullivan, Hongying Li, Aleksei A. Chmura, Alice Latinne, Christian Lange, Tammie O'Rourke, Sarah Olson, Lucy Keatts, A. Patricia Mendoza, Alberto Perez, Cátia Dejuste de Paula, Dawn Zimmerman, Marc Valitutto, Matthew LeBreton, David McIver, Ariful Islam, Veasna Duong, Moctar Mouiche, Zhengli Shi, Prime Mulembakani, Charles Kumakamba, Mohamed Ali, Nigatu Kebede, Ubald Tamoufe, Samuel Bel-Nono, Alpha Camara, Joko Pamungkas, Kalpy J. Coulibaly, Ehab Abu-Basha, Joseph Kamau, Soubanh Silithammavong, James Desmond, Tom Hughes, Enkhtuvshin Shiilegdamba, Ohnmar Aung, Dibesh Karmacharya, Julius Nziza, Daouda Ndiaye, Aiah Gbakima, Zikankuba Sajali, Supaporn Wacharapluesadee, Erika Alandia Robles, Benard Ssebide, Gerardo Suzán, Luis F. Aguirre, Monica R. Solorio, Tapan N. Dhole, Nguyen T. T. Nga, Peta L. Hitchens, Damien O. Joly, Karen Saylors, Amanda Fine, Suzan Murray, William B. Karesh, Peter Daszak, Jonna A. K. Mazet, Cátia Dejuste de Paula, and Christine K. Johnson. Predicting the potential for zoonotic transmission and host associations for novel viruses. *Communications Biology*, 5(1):1–10, August 2022. ISSN 2399-3642. doi: 10.1038/s42003-022-03797-9. URL https://www.nature.com/articles/s42003-022-03797-9.
